# METTL3 facilitates hepatic fibrosis progression via m^6^A-YTHDF2 dependent silencing of GPR161

**DOI:** 10.1101/2021.12.15.472749

**Authors:** Xue-yin Pan, Yi-hui Bi, Miao Cheng, Zhen-zhen Qian, Ling Wang, Hong-mei You, Lei Liu, Zhen-hua Zhang, Xiao-ming Meng, Cheng Huang, Jun Li

**Author notes:** These authors contributed equally to this work (Xue-yin Pan, Yi-hui Bi). Correspondence authors: Jun Li, M.D., Ph.D., School of Pharmacy, Anhui Medical University, Hefei, 230032, China; Cheng Huang, M.D., Ph.D., School of Pharmacy, Anhui Medical University, Hefei, 230032, China.

## Abstract

Hepatic fibrosis (HF) is a very common condition seen in millions of patients with various liver diseases. N^6^-methyladenosine (m^6^A) plays critical roles in various biological and pathological processes. However, the role of m^6^A and its main methyltransferase METTL3 in HF remains obscure. Here, we reported that METTL3 expression was elevated in HSCs from CCl4 induced fibrotic liver. METTL3 knockdown in HSCs mediated by recombinant adeno-associated-virus serotype 9 packed short hairpin RNA against METTL3 alleviated liver injury and fibrosis compared to empty carrier group. Mechanistically, the decreased liver fibrosis in CCl4-treated HSC-specific *METTL3* knockdown mice was due to the increased GPR161 that is a suppressor of Hedgehog pathway, a well-known pathway to activate in liver injury and regeneration. As expect, GPR161 transferred into HSCs alleviated liver fibrosis and HSC activation. Forced GPR161 expression inhibited Gli3 activated form nuclear accumulation and subsequently suppressed fibrosis-associate gene expression. Conclusion, HSC-specific deletion of METTL3 inhibits liver fibrosis via elevated GPR161 expression, which subsequently suppressed Hedgehog pathway activation and fibrosis-associated genes expression, providing novel therapeutic targets for HF therapy.

**Synopsis:** 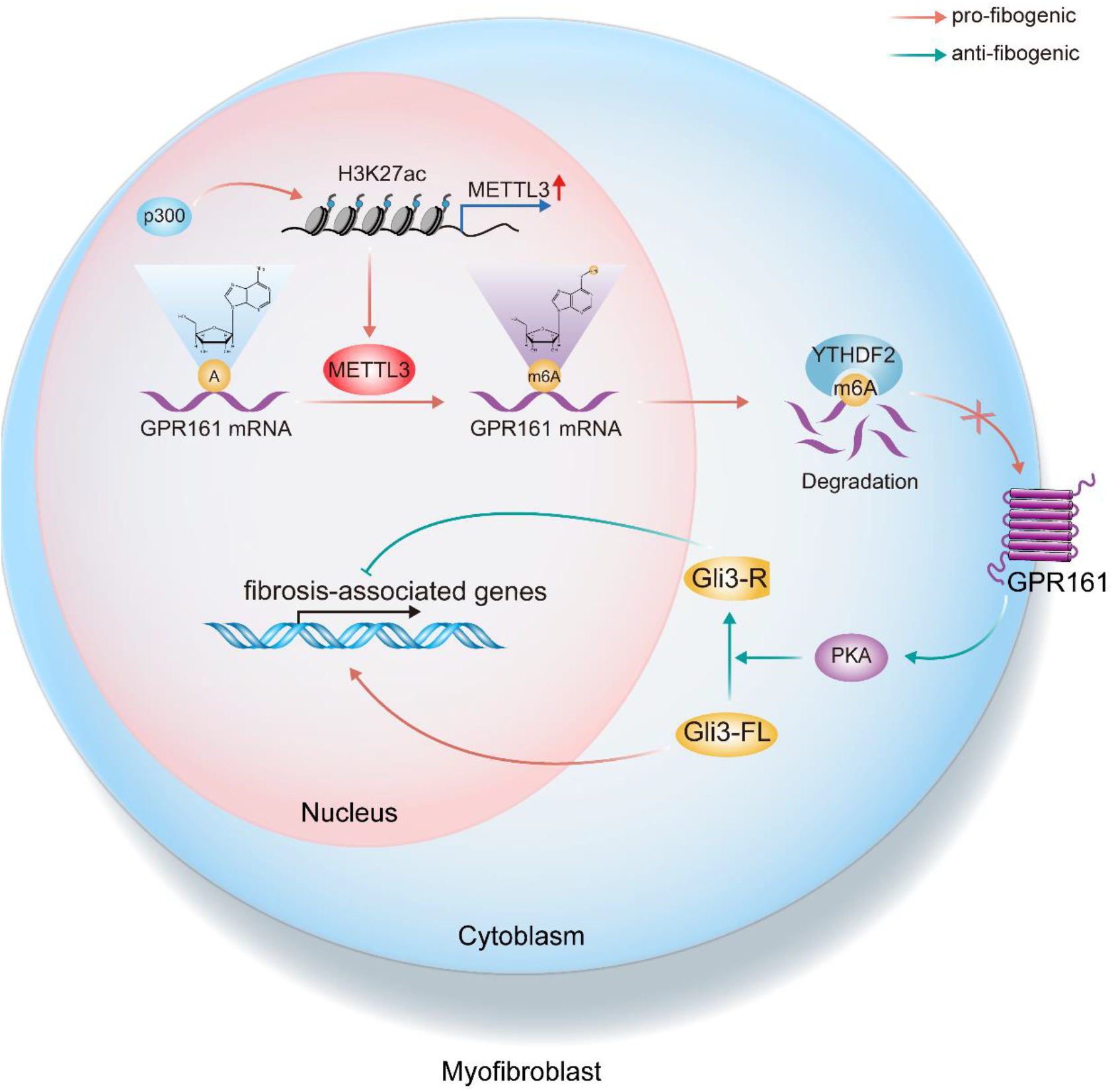

The role of RNA m^6^A methyltransferase METTL3 in the progression of hepatic fibrosis is unclear. Here, we reveal that suppresses m^6^A modification by depletion of METTL3 in Hepatic Stellate Cells (HSCs), stabilizes the GPR161 transcripts, that involved in suppressing Hedgehog signaling pathway, through YTHDF2, thereby alleviating the fibrosis feature of CCl4-induced liver injury.

- Disruption of m^6^A methyltransferases METTL3 in HSCs alleviates liver fibrosis feature.
- METTL3 depletion in liver HSCs suppresses its activation by suppressing Hedgehog signaling pathway through stabilizing GPR161 transcripts in an m^6^A-YTHDF2 dependent manner.
- GPR161 activates PKA that mediates phosphorylation and degradation of full-length Gli3 (Gli3-FL) protein, then the short Gli3 (Gli3-R) translocated into the nucleus and suppresses the fibrosis associated gene transcription.
- The elevated METTL3 expression is attribute to its promoter H3K27ac modification.

## Introduction

Hepatic fibrosis (HF) is a considerable health concern, which is characterized by the increasing extracellular matrix (ECM) components deposition and fibrous scar formation (Bataller & Brenner, 2005, Kisseleva & Brenner, 2020, Tsuchida & Friedman, 2017). Without effective treatment, HF inevitably progresses into liver cirrhosis, liver failure, and/or even hepatocellular carcinoma (HCC), which accounts for million deaths annually worldwide (Asrani, Devarbhavi et al., 2019). Upon liver insult, HSCs trans-differentiate into myofibroblast, participating in the wound healing response of the liver. As the major source of ECM, activated HSCs is a central driver of HF progression (Tsuchida & Friedman, 2017). Understanding the underlying mechanism of HSC activation is therefore indispensable to define novel and more efficient targets of antifibrotic therapy to lower the incidence, morbidity and mortality of the people suffering from chronic liver disease.

N^6^-methyladenosine (m^6^A) is the mostly prevalent internal mRNA modification which is mediated by the main methyltransferase complex METTL3/METTL14/WTAP, and removed by m^6^A demethylases FTO and ALKBH5 (Wei, Gershowitz et al., 1975, Zaccara, Ries et al., 2019). The dynamic m^6^A modification sites on mRNAs are recognized by different “readers”, including YTHDF1/2/3, YTHDC1/2, which are involved in regulating RNA splicing (Xiao, Adhikari et al., 2016), nuclear export (Roundtree, Luo et al., 2017), stability (Wang, Lu et al., 2014), translation (Wang, Zhao et al., 2015), decay (Park, Ha et al., 2019), and structural switches (Liu, Dai et al., 2015). Studies have indicated that m^6^A modification sites preferably occur in the consensus motif “DRm^6^ACH” (D = A, G or U; R = G or A; H = A, C or U) are enriched in the 3′ UTR and CDS, particularly around the stop codon (Dominissini, Moshitch-Moshkovitz et al., 2012, Linder, Grozhik et al., 2015, Meyer, Saletore et al., 2012). Numerous studies have suggested that altered expression of key genes, which are sensitive to m^6^A regulators, may lead to significant phenotypic changes (Mendel, Chen et al., 2018, Shen, Sheng et al., 2020, Shi, Zhang et al., 2018, Weng, Huang et al., 2018, Zhong, Yu et al., 2018, Zhou, Wan et al., 2015). However, whether m^6^A modification involved in regulating HF progression remains obscure.

Therefore, in the present study, to explore the role of m^6^A methylation in the development of HF, we targeted METTL3, a key component of m^6^A methylase, in HSCs. The results of our study revealed that METTL3 regulated GPR161 silencing and played a critical role in HF progression and HSC activation. Thus, an analysis of the mechanism underlying HSC activation is warranted, because it may provide insights into the future of clinical cancer therapies and drug development.

## Results

### Elevated METTL3 expression in fibrotic livers

To investigated the role of m^6^A modification in HF, we employed CCl4-induced chronic liver injury to generate HF model in C57BL/6J mice (**Fig EV1A-E**). Then, we assessed the m^6^A methylation contents on RNAs from CCl4-treated fibrotic livers by the m^6^A RNA methylation quantification kit. As shown in **Fig 1A**, the m^6^A levels were higher in RNAs from CCl4 treated livers than from Oil treated livers. Subsequently, we evaluated the expression changes of m^6^A modulators in fibrotic livers using RT-qPCR analysis. As shown in **Fig 1B**, the expression of METTL3 and WTAP was significantly elevated in the fibrotic livers. As METTL3 is the only catalytic subunit of m^6^A methylase complex, we selected METTL3 for the further study. Next, we verified the elevated METTL3 expression in fibrotic livers using western blot and IHC assay (**Fig 1C** and **EV1C**). The results in BDL model and human liver cirrhosis tissues were consistent with CCl4 induced fibrosis liver (**Fig EV1D-J**). Collectively, these data illustrated that METTL3 expression was significantly increased in the fibrotic livers. Then we performed dual-immunofluorescence assay. The co-localization of METTL3 and α-SMA revealed that METTL3 was expressed in HSCs (**Fig 1D**). Moreover, METTL3 was significantly elevated in primary HSCs from fibrotic livers (**Fig 1E**). Considered together, these data revealed that METTL3 was significantly elevated in fibrotic livers, especially in HSCs.

**Figure 1.**
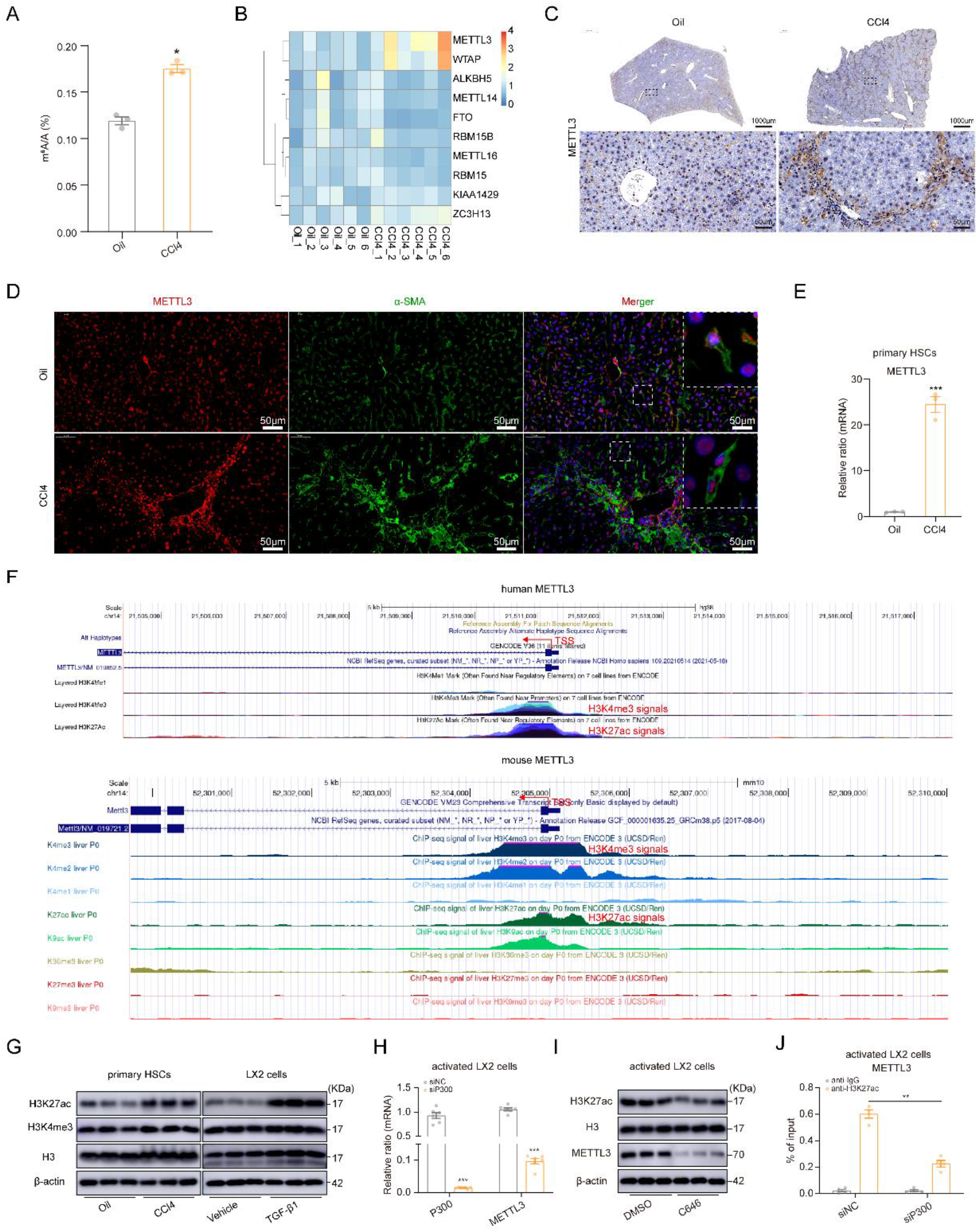
P300 mediated H3K27ac modification activates METTL3 transcription in the HF model. (A) The global m^6^A RNA methylation levels in liver tissues from CCl4- or Oil-treated mice. (B) The m^6^A regulator expression in liver tissues from CCl4- or Oil-treated mice. (C) Liver tissues were subject to METTL3 immunostaining. (D) Immunofluorescence staining in liver tissues from CCl4- or Oil-treated mice. Anti-α-SMA (green), anti-METTL3 (red), and DAPI (blue) were used. Image of DAPI (blue) staining are not shown separately. (E) METTL3 expression in primary HSCs from CCl4- or Oil-treated mice. (F) Data from the UCSC (http://genome.ucsc.edu/) exhibited high enrichment of H3K4me3 and H3K27ac signals in the promoter of METTL3. (G) H3K4me3 and H3K27ac expression in primary HSCs from Oil or CCl4-treated mice, or in LX2 cells cultured with Vehicle or TGF-β1 (40 ng/mL). (H) RT-qPCR analysis for P300 and METTL3 in P300 silencing activated LX2 cells. (I) Western blot analysis for H3K27ac and METTL3 in activated LX2 cells treated with DMSO or C646 (20 µM) for 24 hours. (J) ChIP-qPCR assay was used to detect the enrichment of H3K27ac signals at the promoter of METTL3 in control (siNC) or P300 deficiency (siP300) activated LX2 cells. IgG was performed as the control. Scale bar are shown in the Figure. The data are present as mean ± SEM of at least three independent replicates. * *p* < 0.05; ** *p* < 0.01; *** *p* < 0.001. Source data are available online for this figure.

### Elevated METTL3 expression was attribute to P300-mediated H3K27 acetylation in the HSC from fibrotic liver

To explore whether the histone modification involved in regulating METTL3 expression, we analyzed the histone modification of METTL3 promoter via the UCSC Genome Browser. As presented in **Fig 1F**, abundant H3K4me3 and H3K27ac signals, which was associated with transcription activation, were enriched in the promoter region of METTL3 both in human and mouse species. The protein expression of H3K27ac was significantly elevated in primary HSCs from fibrotic livers and LX2 cells activated with TGF-β1, while H3K4me3 levels was consistent in the two groups (**Fig 1G**), suggesting that H3K27ac might be implicated in regulating METTL3 expression in HSCs.

H3K27ac is catalyzed by the histone acetyltransferase complex P300/CBP, P300 expression was elevated in livers of mice following injection of CCl4, and P300 nuclear accumulation is necessary for HSC activation (Dou, Liu et al., 2018). To investigate whether the enriched H3K27ac signals on METTL3 promoter was mediated by P300 in HSCs, we suppressed P300 expression and activation using siRNA against P300 (**Fig S1K**) or C646 (an P300 inhibitor). P300 inhibition significantly suppressed H3K27ac and METTL3 expression in activated LX2 cells (**Fig 1H-I** and **EV1L**). Moreover, chromatin immunoprecipitation (ChIP) assay indicated that P300 suppression substantially decreased the enrichment of H3K27ac signals on METTL3 promoter (**Fig 1J**). Considering together, these data illustrated that P300 mediated H3K27ac might partly account for the upregulation of METTL3 expression in the HF model.

### HSC-specific METTL3 silencing attenuates the pathology of liver fibrosis

To explore the role of METTL3 in HF HSCs, we disrupted METTL3 expression specially in HSCs using Ad-Lrat-shM3-Luc. The bioluminescence image of mice at indicated time point showed the enrichment of virus in the liver (**Fig 2A** and **EV2A**). Furthermore, the immunofluorescence assay revealed that METTL3 expression was significantly decreased in the fibrotic areas of liver tissues (**Fig 2B**). In addition, METTL3 expression was specifically suppressed in primary HSCs from Ad-Lrat-shM3-Luc transfected mice treated with CCl4 (**Fig 2C**). METTL3 silencing in HSCs significantly suppressed the inflammatory cells infiltration, ECM deposition, serum ALT, and AST levels in fibrotic mouse model (**Fig 2D-E**). Furthermore, the expression of fibrosis markers was significantly decreased in the primary HSCs from the Ad-Lrat-shM3-Luc transfected mice (**Fig 2F-G**). Collectively, these results suggested that METTL3 silencing in HSCs alleviates the pathology of HF.

**Figure 2.**
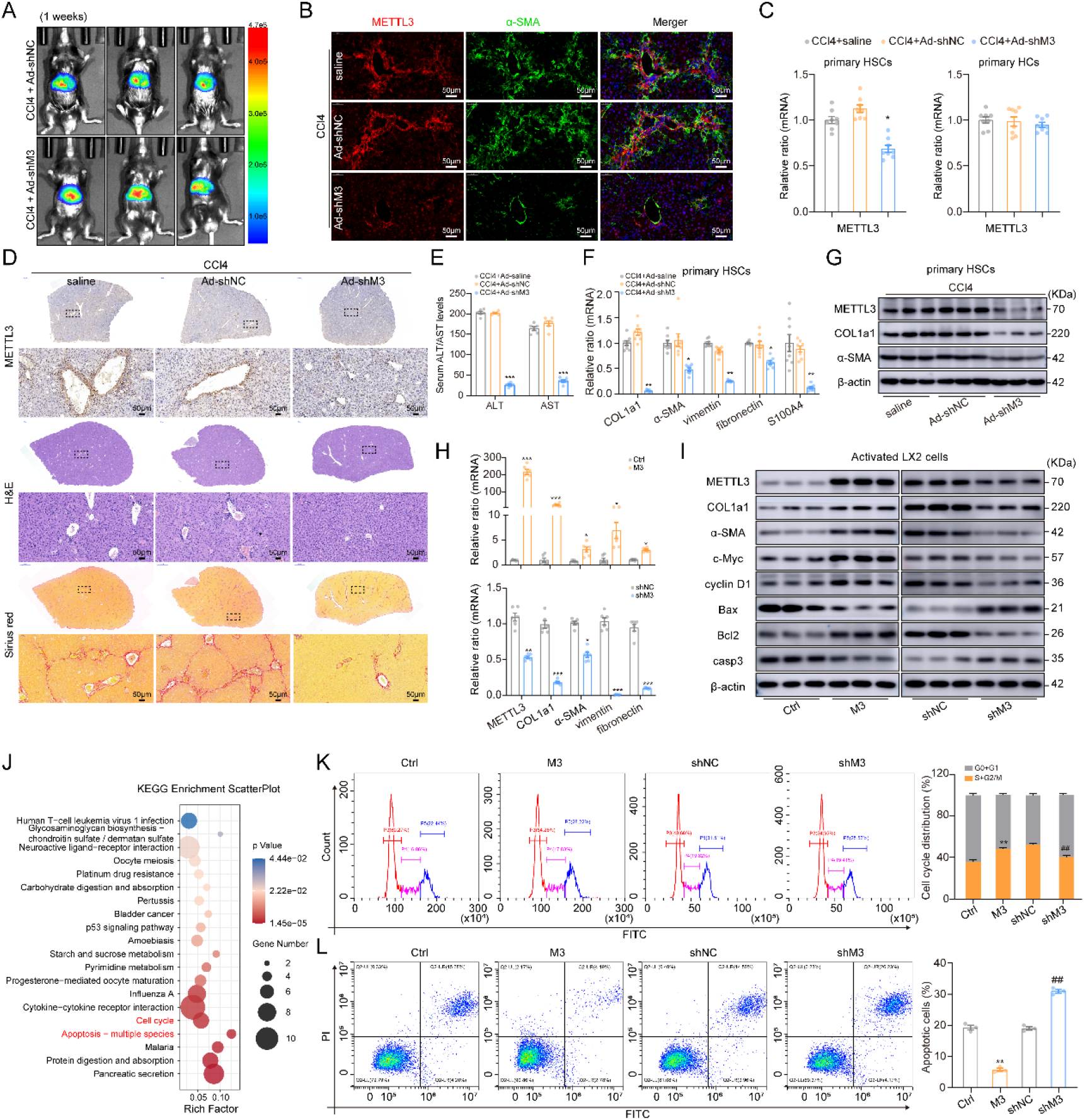
HSC-specific METTL3 silencing alleviates HF pathology *in vivo* and *in vitro*. (A) Bioluminescence imaging of Ad-Lrat-shM3-Luc (Ad-shM3) and Ad-Lrat-shNC-Luc (Ad-shNC) transfected mice (one week after virus infection). (B) Immunofluorescence in liver tissue from Ad-shNC and Ad-shM3 transfected mice, and saline-treated mice. Anti-α-SMA (green), anti-METTL3 (red), and DAPI (blue) were used. Image of DAPI (blue) staining are not shown separately. (C) METTL3 expression were detected in primary HSCs and primary HCs from Ad-shNC and Ad-shM3 transfected mice, and saline-treated mice. (D) Representative images of METTL3 immunostaining, H&E staining, and Sirius red staining of liver tissue sections are shown. (E) Serum ALT and AST levels were measured. (F) RT-qPCR analyses of *COL1a1*, *α-SMA*, *vimentin*, *fibronectin*, and *S100A4* expression, and (G) western blot analyses of METTL3, COL1a1, and α-SMA expression in primary HSCs from Ad-shNC and Ad-shM3 transfected mice, and saline-treated mice. (H) RT-qPCR analyses of *METTL3*, *COL1a1*, *α-SMA*, *vimentin*, and *fibronectin* were measured in activated LX2 cells with stable METTL3 overexpressed (upper panel) or stable METTL3 suppression (lower panel). (I) Western blot analysis for METTL3, COL1a1, α-SMA, c-Myc, cyclin D1, Bax, Bcl2, and Caspase 3 expression in stable METTL3 overexpressing (left panel) and silencing (right panel) in activated LX2 cells. (J) KEGG enrichment analysis of different expressed genes (|log2(fc)| ≥ 1; *p* < 0.05) in METTL3 silencing (siM3) compared with negative control (siNC) transfected primary HSCs. (K-L) The effect of loss- and gain-of expression of METTL3 on the cell cycle distribution and the percentage of apoptosis cells in activated LX2 cells was detected by flow cytometry. Scale bar = 50 μm. The data are present as mean ± SEM of at least three independent replicates. * *p* < 0.05; ** *p* < 0.01; *** *p* < 0.001; ^##^ *p* < 0.01. Source data are available online for this figure.

### METTL3 regulates HSC proliferation, migration, and apoptosis

To elucidate the role of METTL3 in HSCs, both gain- and loss-of-function studies were conducted in human LX2 cells. We retrovirally transduced METTL3 or shMETTL3 and corresponding control lentivirus into LX2 cells and selected individual stable clones using puromycin (1 mg/mL). The eGFP signals indicated successful transfection of the lentivirus (**Fig EV2B**). *METTL3*-overexpressing stable cell lines exhibited a 14- to 15-fold increase in *METTL3* mRNA levels, while *METTL3*-silencing stable cell lines exhibited a 9- to 10-fold decrease (**Fig EV2C**), accompanied by a noticeable increase or decrease in the m^6^A modification levels, respectively (**Fig EV2D-E**) compared with those of the control cell lines. Increase in METTL3 knock in, and decrease in METTL3 knock down were observed for fibrosis markers in activated LX2 cells (**Fig 2H-I**). The function of METTL3 in primary HSCs was consistent with results in LX2 cells (**Fig EV2F-H**). These results indicated that METTL3 plays critical role in regulating HSC activation.

To gain an overview of the global role of METTL3 in HSCs, we performed RNA sequencing (RNA-Seq) in METTL3 silencing primary HSCs treated with TGF-β1 for 24 hours. The data revealed that 111 upregulated and 180 downregulated genes were significantly altered in RNA-seq analysis from primary HSCs with knock down METTL3 compared to control (**Dataset EV1**). The KEGG pathway analysis revealed that METTL3 silencing affect a list of genes associated with cell cycle and apoptosis pathway (**Fig 2J**). Subsequently, we investigate the role of METTL3 in regulating cell cycle and apoptosis of HSCs. As shown in **Fig 2I**, c-Myc and cyclin D1 protein levels were increased in METTL3 overexpression group in activated LX2 cells, but decreased in METTL3 ablation group. The ratio of Bax/Bcl2 and Caspase3 were decreased in the METTL3 overexpression group in activated LX2 cells, but increased in METTL3 silencing group (**Fig 2I**). Furthermore, the flow cytometry analyses revealed that METTL3 overexpression enhanced the cell cycle of HSC distributed in the S and G_2_/M phase in activated LX2 cells, while METTL3 silencing inhibited it (**Fig 2K**). A decrease in the proportion of apoptotic cells in the METTL3 overexpression group in activated LX2 cells, and an increase in the METTL3 silencing group was observed (**Fig 2L**). EdU assay revealed that METTL3 knock in promoted the proliferation of activated LX2 cells, while METTL3 knock out suppressed it (**Fig EV2I-J**). Transwell assay and scratch wound healing assay showing that METTL3 overexpression promoted the migration of activated LX2 cells, while METTL3 depletion suppressed it in activated LX2 cells (**Fig EV2K-L**). Considering together, these data indicated that METTL3 played a pivotal role in promoting HSC proliferation and migration, and in inhibiting HSC apoptosis.

### Variation of m^6^A-regulated genes in the HF mouse model

To comprehensively profile genes with m^6^A modification in HF process, we performed MeRIP-Seq in liver tissues after CCl4 induced fibrosis. The scheme of MeRIP-Seq and RNA-Seq processes was shown in **Fig 3A**, and the classificational analysis of differentially expressed RNAs was shown in **Fig 3B**. Moreover, 93 to 174 million reads were generated from each MeRIP-Seq or RNA-Seq (also served as the input control for the corresponding MeRIP-Seq library) (**Table EV1**). A total of 1 607 genes were upregulated, while 398 were downregulated in the CCl4-induced fibrotic liver, compared to the Oil control (**Fig EV3A**) (**Dataset EV2**). MeRIP-Seq analysis identified 48 725 and 51 222 from 15 725 and 16 528 m^6^A-modified genes in Oil and CCl4 group, respectively. Peak distribution analysis revealed strong enrichment of m^6^A peaks near the stop codon (**Fig EV3B**) as previously described (Kate D. Meyer, Yogesh Saletore et al., 2012). The m^6^A consensus motif GGAC was identified in fibrotic livers (**Fig EV3C**). GO analysis revealed that m^6^A-methylated mRNAs are involved in critical cellular processes, such as cell differentiation, cell cycle, DNA damage, apoptotic process, transcription, suggesting that m^6^A modification plays pivotal role in HF.

**Figure 3.**
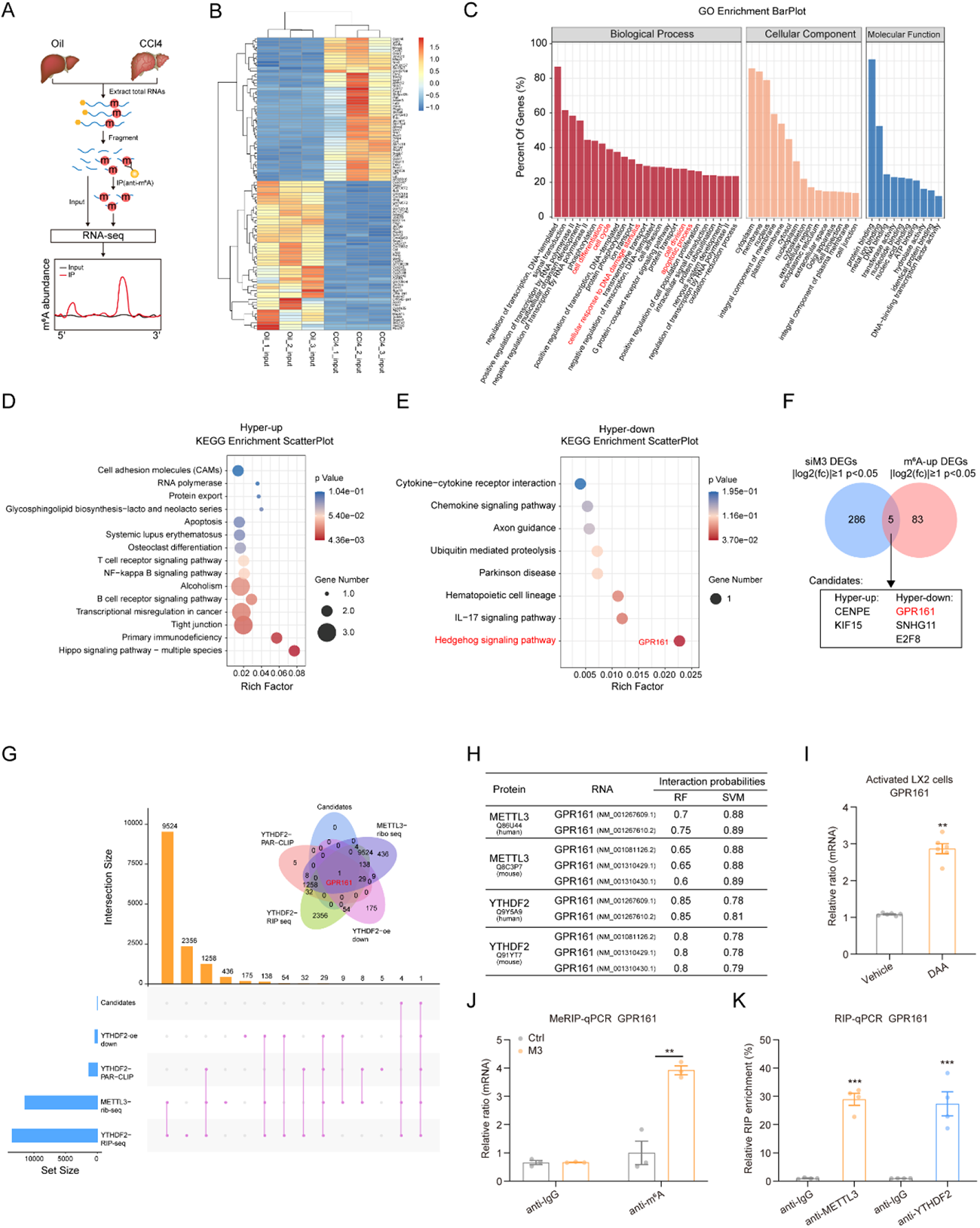
m^6^A methylation underlies the effects of METTL3. (A) Schematic of the MeRIP-Seq protocol used to identify the m^6^A methylation landscape following fibrosis mouse model establishment. (B) Heatmap of differentially expressed genes (DEGs) identified by RNA-Seq. The blue and red color indicate downregulation and upregulation of expression levels, respectively (|log2(fc)| ≥ 1; *p* < 0.05). (C) GO enrichment analysis of genes with m^6^A peak deposition. (D-E) KEGG enrichment analysis of Hyper-up genes and Hyper-down genes in livers from CCl4-treated mice compared with from Oil-treated mice. (F) Venn diagram shows the numbers of DEGs that also have the m^6^A mark in METTL3 silencing group. (G) Overlapping genes in Candidates, YTHDF2-PAR-CLIP (GSE49339), YTHDF2-RIP (GSE49339), YTHDF2-oe downregulated gene (PMID: 31735169) and METTL3-Ribosome (GSE63591) are shown. (H) The interactions established between GPR161 mRNA and METTL3 or YTHDF2 protein in mouse and human were predicted using RNA-Protein Interaction Prediction (RPISeq) (http://pridb.gdcb.iastate.edu/RPISeq/). (I) GPR161 expression in activated LX2 cells treated with 3-deazaadenosine (DAA) or Vehicle (DMSO) for 24 hours. (J) The m^6^A methylation levels of GPR161 mRNA were determined by MeRIP-qPCR. (K) Binding of METTL3 or YTHDF2 protein with GPR161 mRNA was analyzed via an RIP-qPCR assay. The data are present as mean ± SEM of at least three independent replicates. ** *p* < 0.01; *** *p* < 0.001. Source data are available online for this figure.

A total of 691 and 286 m^6^A peaks showed a significant increase and decrease (|log_2_(fc)| ≥ 1; *p* < 0.05), respectively, in abundance, in the CCl4-treated group compared to the Oil group. Thus, these were termed hyper- and hypo-methylated m^6^A peaks, respectively. Through analysis of the RNA-Seq data, we identified 691 hypermethylated m^6^A peaks of the mRNA transcripts that were significantly (|log_2_(fc)| ≥ 1; *p* < 0.05) upregulated (81; Hyper-up) or downregulated (13; Hyper-down) in the CCl4-treated group relative to the Oil group (**Fig EV3D**) (**Dataset EV2**). Consistently, we identified 286 hypomethylated m^6^A peaks of the mRNA transcripts that were significantly (|log_2_(fc)| ≥ 1; *p* < 0.05) upregulated (31; Hypo-up) or downregulated (6; Hypo-down) in the CCl4-treated group relative to the Oil group (**Fig EV3D**). Subsequently, KEGG pathway show that Hyper-up genes involved in regulating Hippo pathway (**Fig 3D**), and Hyper-down genes involved in modulating Hedgehog pathway (**Fig 3E**), which has been reported play pivotal role in regulating HSC activation and liver fibrosis (Du, Hyun et al., 2018). Collectively, these data illustrated that m^6^A modification involved in regulating HF process.

### METTL3 regulated GPR161 expression in an m^6^A-YTHDF2 dependent manner

To find the genes which were regulated by METTL3 in HF and in HSC trans-differentiation process, we overlapped the hypermethylated genes in CCl4 group with genes differently expressed in siMETTL3 transfected HSCs (|log_2_(fc)| ≥ 1; *p* < 0.05) (**Fig 3F**) (**Table EV2**). Then, we focused on GPR161, that is a conserved vertebrate G-protein-coupled receptor and involved in negatively regulating Hedgehog pathway. We noticed that the m^6^A methylation levels of GPR161 mRNA was elevated but the mRNA levels were decreased. Previous studies illustrated that YTHDF2 is the major m^6^A reader which mediated degradation of its target transcripts (Park et al., 2019, Wang et al., 2014). Then, we hypothesized that the decreased expression of GPR161 mRNA might be attribute to YTHDF2-mediated mRNA degradation. To explore this hypothesis, we overlapped the Candidates in our study with genes in YTHDF2-PAR-CLIP-Seq; YTHDF2-RIP-Seq; decreased genes in YTHDF2-oe group; and METTL3-Ribo-Seq. As shown in **Fig 3G**, the only overlapped gene was GPR161. Subsequently, we reasoned the interaction between *GPR161* mRNA with METTL3 protein and YTHDF2 protein using the RNA-protein interaction prediction (RPISeq) database, based on the random forest (RF) model or support vector machine (SVM), and probabilities > 0.5 were considered “positive” (**Fig 3H**). In addition, treatment of activated LX2 cells with 3-deazaadenosine (3-DA), a global methylation inhibitor, substantially increased *GPR161* mRNA levels, suggesting that GPR161 expression might be regulated by RNA methylation (**Fig 3I**). Furthermore, MeRIP-qPCR assay indicated that *GPR161* mRNA was methylated (**Fig 3J**), and RIP-qPCR assay suggested that *GPR161* mRNA established interaction with the METTL3 and YTHDF2 protein in activated LX2 cells (**Fig 3K**). Moreover, METTL3 overexpression inhibited, while METTL3 silencing elevated GPR161 protein and mRNA expression *in vivo* and *in vitro* (**Fig 3E-H**). In addition, knockdown METTL3 extended the half-life of the *GPR161* transcript (**Fig EV3I**). Next, we knockdown YTHDF2 expression using lentivirus-packed shYTHDF2 (**Fig EV3J**). YTHDF2 silencing elevated GPR161 expression at the protein and mRNA levels in activated LX2 cells (**Fig EV3K-L**). Considered together, these data suggested that the expression of GPR161 was regulated by METTL3 in an m^6^A-YTHDF2-dependent manner.

### METTL3 regulates GPR161 expression relies on its methyltransferase ability

Through analyzing the MeRIP-Seq data, we found that *GPR161* transcripts have several significant m^6^A peaks on CDS and 3′ UTR region (**Fig 4A**). Then, we predicted the high confidence m^6^A sites in *GPR161* mRNA by SRAMP (**Fig 4B**), and these highly confident m^6^A sites on CDS and 3′ UTR region of *GPR161* mRNA were presented in **Fig 4C**. To address the potential role of m^6^A modification on the CDS and 3′ UTR of *GPR161* mRNA, we constructed luciferase reporter plasmids, which contained the wild-type or mutant CDS or 3′ UTR of *GPR161* (“A” in m^6^A motif was replaced by “G”) (**Fig 4D**). Given that point mutations (C294A and C326A) located in the CCCH motif of METTL3, full-length complexes lose their m^6^A methylase activity (Wang, Doxtader et al., 2016), and the amino acid sequence of METTL3 in human and mouse species were conserved (96.9%), as well as the point of C294 and C326 (**Fig 4E**). Therefore, we constructed point mutation (C294A and C326A) plasmid of METTL3 in mouse species. Compared to the control plasmid, METTL3 overexpression substantially elevated luciferase activity of the individual reporter constructs harboring wild-type *GPR161* CDS and 3′ UTR that contained intact m^6^A sites (**Fig 4F-G**). However, METTL3 overexpression did not exert a significant effect on the luciferase activity in the reporter plasmids harboring mutant m^6^A motifs (**Fig 4F-G**). METTL3 mutant plasmid did not influence on the luciferase activity of report harboring wild-type *GPR161* CDS and 3′ UTR that contained intact m^6^A sites (**Fig 4F-G**). Considered together, our data indicated that METTL3 suppressed GPR161 expression depends on its m^6^A methyltransferase activity.

**Figure 4.**
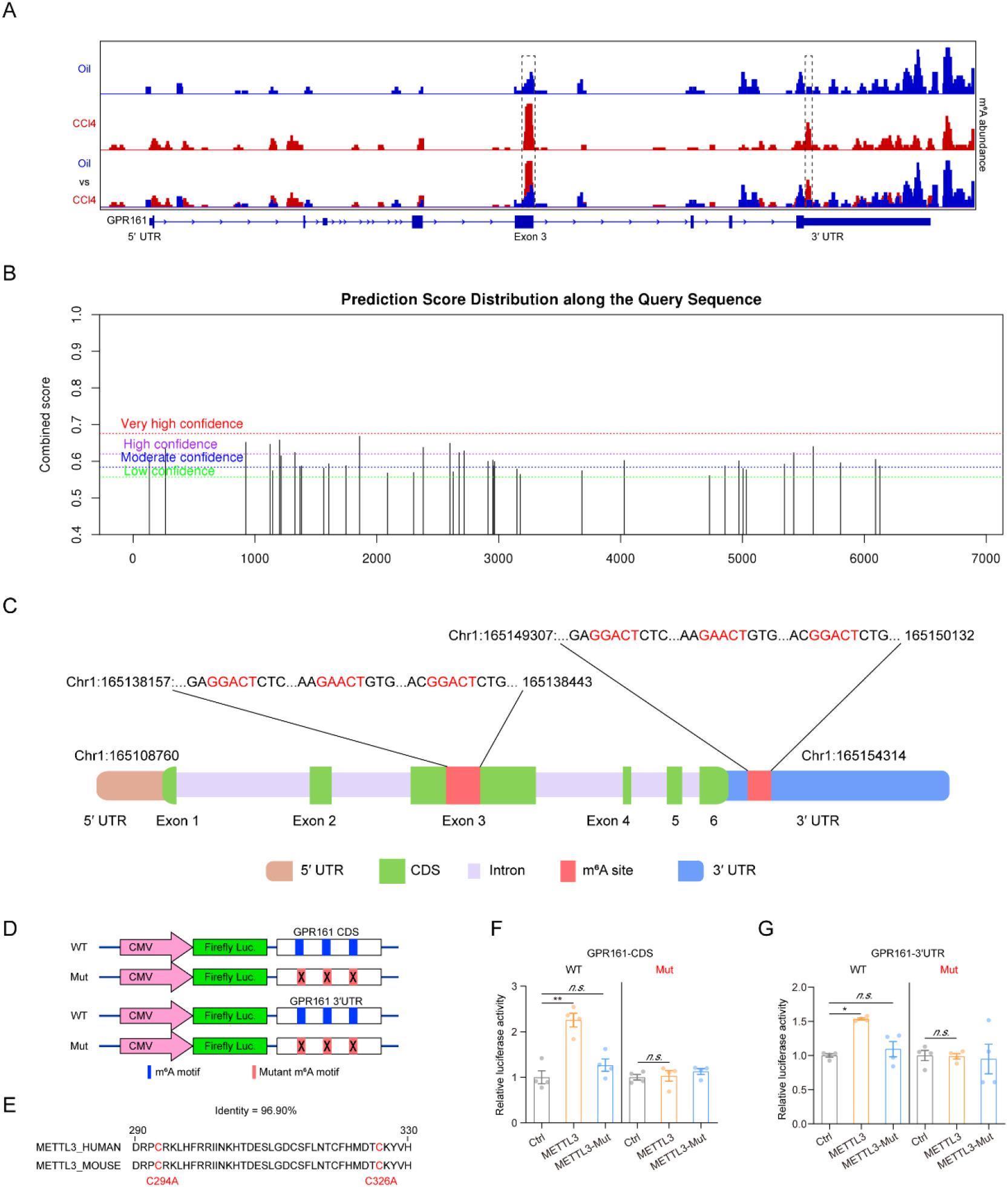
METTL3-mediated m^6^A deposition on *GPR161* mRNA decreases its stability in an m^6^A-YTHDF2 dependent manner. (A) Distributions of the m^6^A peaks on *GPR161* mRNA were visualized using the Integrative Genomics Viewer (IGV). Square indicate elevated m^6^A peaks in the liver tissues from CCl4-treated mice compared with Oil-treated mice. (B) The m^6^A modification sites were predicted using the SRAMP online tool (http://www.cuilab.cn/sramp). (C) Schematic of the positions of the m^6^A modification sites on *GPR161* mRNA CDS and 3′ UTR. (D) Schematic of mutation in GPR161 mRNA CDS and 3′ UTR. (E) Schematic showing the evolutionarily conserved cysteine residue (C294 and C326) of METTL3 in human and mouse. (F-G) Relative luciferase activity of pcDNA3.1-reporter-GPR161-CDS and pcDNA3.1-reporter-GPR161-3′ UTR with either a wild-type or mutant (A-to-G mutant) m^6^A site following co-transfection with control vector (Ctrl), METTL3 (METTL3), or METTL3 mutant (METTL3-mut) into primary HSCs. Firefly luciferase activity was detected and normalized via Renilla luciferase activity. The data are present as mean ± SEM of at least three independent replicates. * *p* < 0.05; ** *p* < 0.01.

### GPR161 overexpression suppresses HF *in vivo* and *in vitro*

GPR161 expression was decreased in activated HSCs (**Fig EV4A-B**). Forced GPR161 expression alleviated *COL1a1* and *α-SMA* expression *in vitro* (**Fig EV4C-E**). To investigate the role of GPR161 *in vivo*, Ad-Lrat-GPR161-eGFP virus was used to specially elevated GPR161 expression in HSCs. The eGFP signals indicated successful infection (**Fig 5A**). The dual-immunofluorescence assay suggested the overlapped GPR161 expression with eGFP (**Fig 5B**). As shown in **Fig 5C**, the GPR161 expression was substantially elevated in primary HSCs, rather than primary HCs, from Ad-Lrat-GPR161-eGFP virus transfected fibrotic livers. Forced GPR161 expression in HSCs associated with lower inflammatory cells infiltration, ECM deposition, serum ALT and AST levels, and fibrosis marker expression in fibrotic livers than in empty virus transfected mice (**Fig 5D-G**). Considered together, these data suggesting that elevated GPR161 expression in HSCs alleviates HF process.

**Figure 5.**
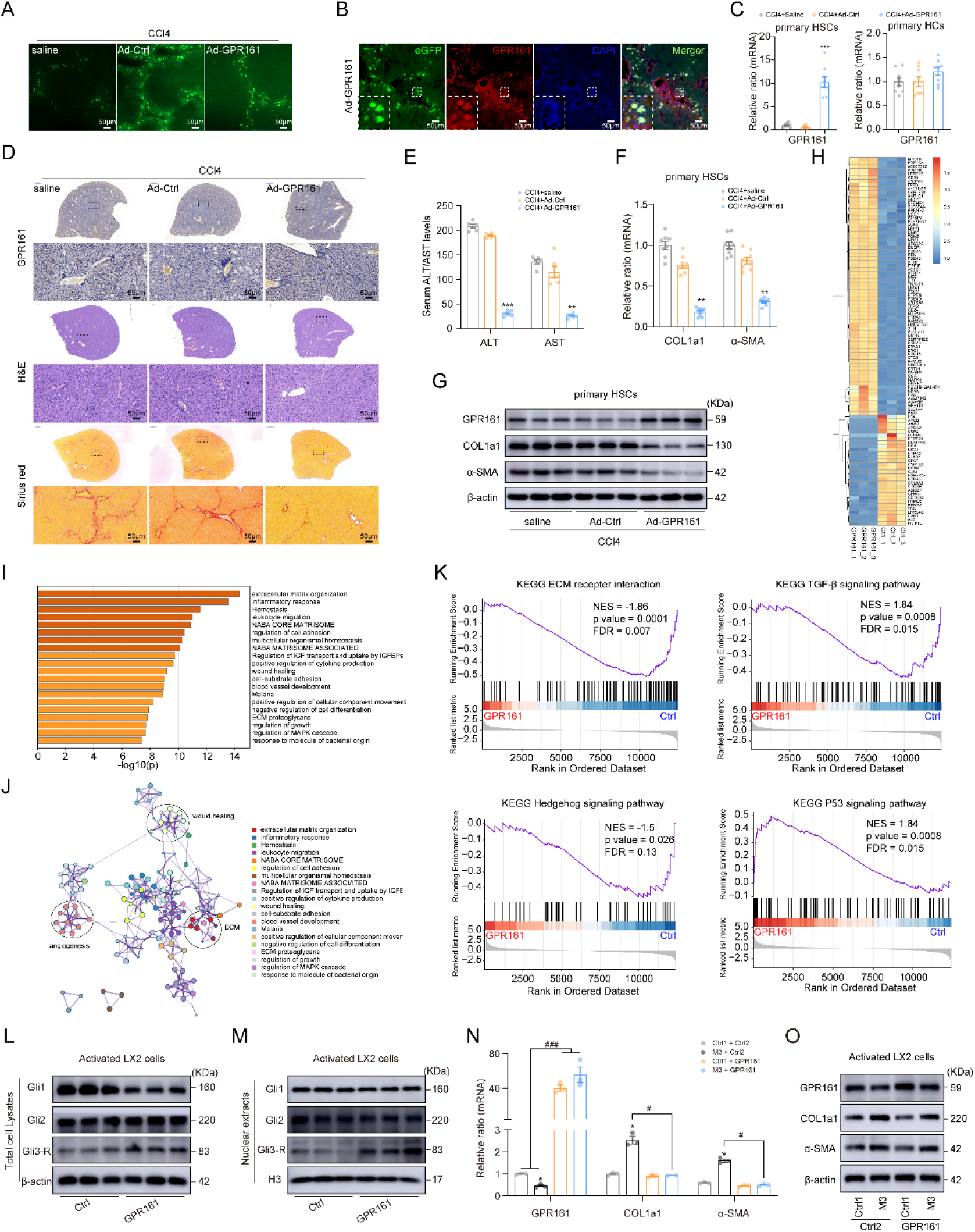
METTL3 regulates HSC proliferation by influencing the GPR161/Gli3 axis. (A) The eGFP signals of liver tissues transfected with Ad-Lrat-GPR161-eGFP (Ad-GPR161) and Ad-Lrat-Ctrl-eGFP (Ad-Ctrl). (B) Immunofluorescence staining for GPR161 in liver tissues from saline treated, Ad-Ctrl or Ad-GPR161 transfected mice following CCl4-induces fibrosis mouse model. The eGFP (green), anti-GPR161 (red), and DAPI (blue) were used. (C) GPR161 expression in primary HSCs and primary HCs from saline treated, Ad-Ctrl or Ad-GPR161 transfected mice following CCl4-treated. (D) Representative pictures of METTL3 immunostaining, H&E staining, and Sirius red staining. (E) Serum ALT and AST levels. (F) RT-qPCR analysis of *COL1a1*, and *α-SMA* in primary HSCs from saline-treated, Ad-Ctrl or Ad-GPR161 transfected mice following CCl4-induced fibrosis. (G) Western blot analysis of GPR161, COL1a1, and α-SMA in primary HSCs from saline-treated, Ad-Ctrl or Ad-GPR161 transfected mice following CCl4-treated. (H) RNA-Seq was performed with RNA extracted from activated LX2 cells with stable elevated GPR161expression. (I) Heatmap of GO enriched terms colored by clusters. (J) Network of enriched terms colored by cluster ID, where nodes that share the same cluster ID are typically close to each other. (K) GSEA plots showing the pathways of DEGs altered by GPR161 was involved in activated LX2 cells. (L-M) Protein levels of Gli1, Gli2, and Gli3 in the total lysates and nuclear extracts were measured by conducting western blot analysis. (N-O) The stable METTL3 overexpressed LX2 cells was transfected with lentivirus packed GPR161 overexpression plasmid. The expression of GPR161, COL1a1, and α-SMA were measured. The data are present as mean ± SEM of at least three independent replicates. * *p* < 0.05; ** *p* < 0.01; *** *p* < 0.001; ^#^ *p* < 0.05; ^###^ *p* < 0.001. Source data are available online for this figure.

### GPR161 regulates HSCs activation and apoptosis

To investigate underlying molecular mechanism, the RNAs isolated from GPR161 overexpressed LX2 cells was used for conducting RNA-Seq. The data revealed that 364 mRNAs were upregulated and 263 genes were downregulated in the LX2 cells transfected with GPR161 (**Fig 5H**) (**Dataset EV3**). The KEGG pathway analysis revealed that GPR161 overexpression affects a list of genes associated with ECM deposition, would healing, and angiogenesis (**Fig 5I-J**), suggesting that GPR161 harbored the capacity to regulate HF progression. Furthermore, gene set enrichment analysis (GSEA) also revealed that genes altered by GPR161 were associated with signal transduction of ECM receptor interaction, TGF-β, Hedgehog, and P53 signaling pathway (**Fig 5K**), supporting the regulatory roles of GPR161 in activation and proliferation of HSCs. GPR161 overexpression inhibited the protein levels of c-Myc, cyclin D1 and Bcl2, but elevated Bax and Caspase 3 expression in activated LX2 cells (**Fig EV4F**). Forced GPR161 expression inhibited HSC migration (**Fig EV4G**) and induced cell cycle arrest in the G_0_/G_1_ phase (**Fig EV4H**), and promoted apoptosis (**Fig EV4I**) in activated LX2 cells. Considering together, these data indicated that GPR161 overexpression could inhibit HSC activation and ECM deposition.

### GPR161 suppressed HSC activation and HF process via inhibiting Hedgehog pathway

Reactivation of Hedgehog pathway, a morphogen that is crucial for embryogenesis, occurs during liver injury and is pivotal in orchestrating liver repair responses. The Hedgehog pathway can be simplified into four main components: the ligand (Hh), the receptor (PTCH1), the signal transducer Smoothened (SMO), and the effector transcription factor (Gli). In the absence of Hh, PTCH1 repressed SMO activity by preventing it entered the primary cilium. Subsequently, the signals transductors Gli proteins will be phosphorylated by several kinases, such as protein kinase A (PKA). When Hh ligand binding with PTCH1, it abrogated the suppression of PTCH1 to SMO. Activate SMO suppressed phosphorylation and degradation of Gli. Full length Gli (Gli-FL) proteins will translocated to the nucleus where it acts as a transcription factor for several target genes (Machado & Diehl, 2018). It has been illustrated that dead hepatocytes could activate HSCs by release Hh ligands (Chung, Moon et al., 2016).

It has been reported that GPR161 involved in negatively regulate Hedgehog pathway through recruiting and activation PKA which mediated Gli3 phosphorylated degradation in repression pattern (Gli3-R) (Mukhopadhyay, Wen et al., 2013). Our GSEA analysis revealed that GPR161 overexpression inhibited Hedgehog pathway activation in activated LX2 cells (**Fig 5K**). Therefore, to investigate the role of GPR161 on Hedgehog pathway in HSCs, we determined the protein levels of Gli1, Gli2 and Gli3 in stable GPR161 overexpression LX2 cells cultured with TGF-β1. As shown in **Fig 5L**, GPR161 overexpression suppressed the activation of Hedgehog pathway, as indicated by decreased Gli1 expression in GPR161 overexpression LX2 cells. Ectopic GPR161 expression decreased Gli3 protein levels in total lysate and nuclear extracts (**Fig 5L-M**). However, the nuclear and total protein levels of Gli2 was unchanged (**Fig 5L-M**). Subsequently, we performed rescue assay to investigate whether METTL3 promoted HF progression through GPR161. As shown in **Fig 5N-O**, forced GPR161 expression decreased COL1a1 and α-SMA levels in stable METTL3 overexpression LX2 cells treated with TGF-β1, both at mRNA and protein levels. Considered together, these data suggested the importance of METTL3/GPR161/Gli3 axis in promoting HF progression.

## Discussion

Various epigenetic modifications, including DNA methylation, histone modification, non-coding RNA regulating, has been proved to contribute to HF progression (Moran-Salvador & Mann, 2017). However, the involved of m^6^A methylation is poorly understand in HF. In the present study, we show that m^6^A modification levels is significantly increased due to the increased METTL3 expression in fibrotic liver. Mechanistically, P300-mediated H3K27ac signals deposition mediated transcription activation of METTL3, which subsequently mediates the m^6^A modification of *GPR161* transcripts, and then the m^6^A reader YTHDF2 directed recognizes and binds the m^6^A sites on *GPR161* transcripts and induces its degradation. Decreased GPR161 levels promotes HF progression through enhancing nuclear accumulation of Gli3 activated form (Gli3-FL), which subsequently promotes profibrogenic gene transcription (**Fig 6**). This is especially important considering the broader function of m^6^A modification and its related modifiers in various biological processes.

**Figure 6.**
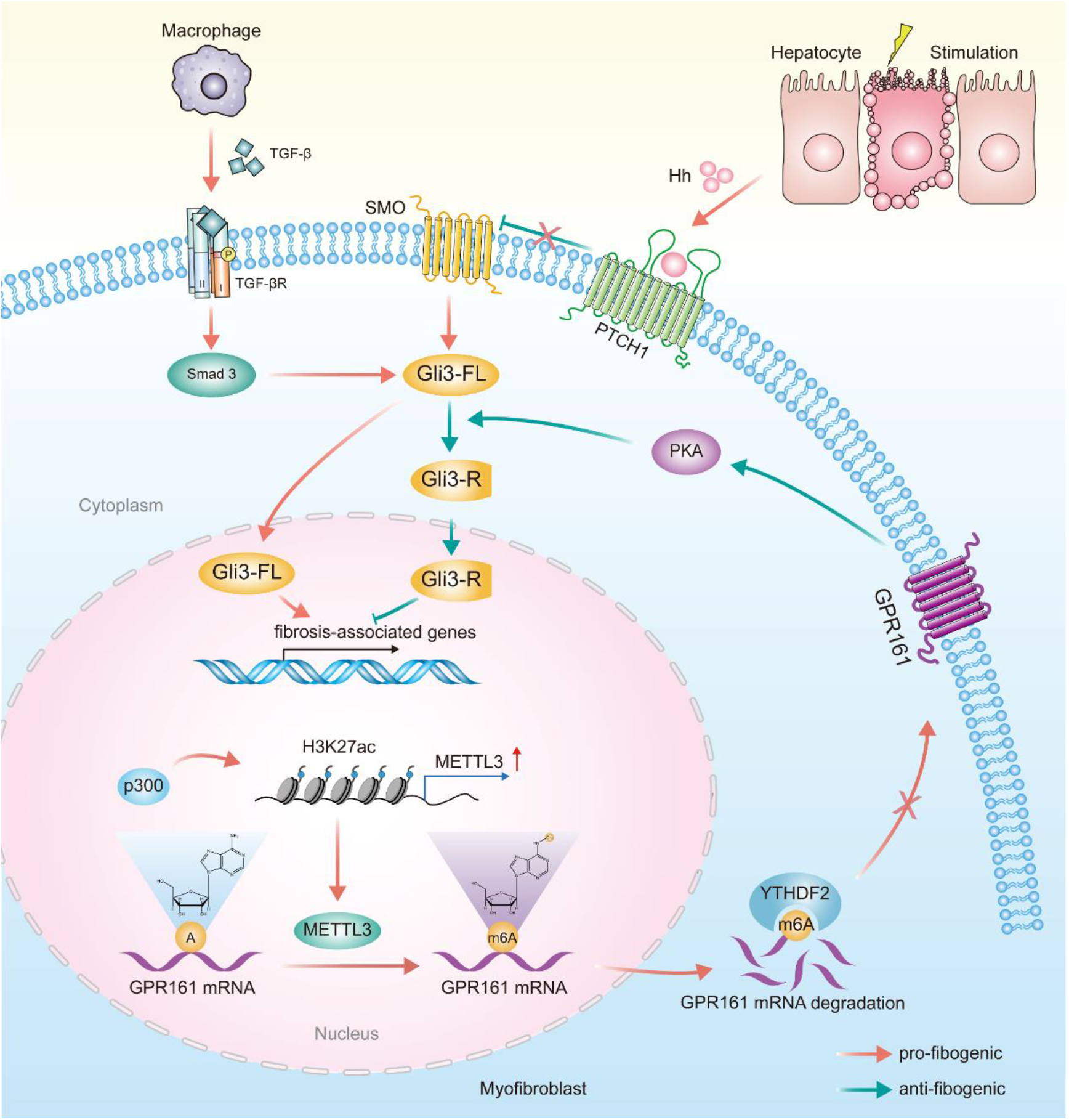
Schematic illustration of METTL3 modulating HF progression and HSC activation. Hepatocyte apoptosis and macrophage activation promote the activation of Hedgehog signaling and nuclear accumulation of full length Gli3 (Gli3-FL), which further induced transcriptome activation of fibrosis-associated genes. Gli3 protein can be phosphorylated and degradation into Gli3 repressor (Gli3-R) by PKA, which was recruited and activated by GPR161. In the model used in present study, P300-mediated H3K27ac induced the transcriptional activation of *METTL3*. Elevated METTL3 expression induced m^6^A methylation deposition on *GPR161* mRNA, which further induced its degradation in an m^6^A-YTHDF2-dependent manner. Decreased GPR161 expression indirectly facilitated the activation of Hedgehog signaling pathway, via abrogating the phosphorylation and degradation of Gli3. Consequently, increased nuclear accumulation of Gli3-FL promoted HF progression and HSC activation by facilitating fibrosis-associated genes expression.

Myofibroblast are the main source of ECM in fibrotic livers. Activated HSCs and activated portal fibroblasts comprise > 90% of the collagen-producing cells, and therefore represent a primary target for antifibrotic therapy (Kisseleva & Brenner, 2020). Our study demonstrates that HSC-specifically METTL3 depletion or GPR161 overexpression alleviates collagen deposition and inflammatory cells infiltration of HF. Our data reveal the crucial role of METTL3 and GPR161 in specifically regulating the activation of HSCs in HF, which establishes the rationale for selectively targeting activated HSCs in HF using METTL3 and GPR161 as a therapeutic target.

H3K27ac is an important mark that associated with increased activation of promoter and enhance elements. This marker can be deposited by both P300 and CREB binding protein. Previous studies have revealed that stiffness induces P300 nuclear accumulation and promotes P300-dependent transcription of α-SMA and CTGF, suggesting a crucial role of P300 in HSC activation (Dou et al., 2018). However, research about the regulation role of H3K27ac in HSC activation and the related gene loci are very limited. Our findings indicate that METTL3 is one of the related loci affected by H3K27ac in HF, and P300-mediated the H3K27ac marks deposition on METTL3 locus. Our study demonstrates that histone modification during HF regulate METTL3 expression.

Through transcriptome-wide RNA-Seq and MeRIP-Seq, we identified a set of potential targets of METTL3, which involved in pathways related to cell cycle, cell proliferation and apoptosis, such as P53, Hippo, NF-KB, Hedgehog pathway. Next, we identified *GPR161* as a direct target of METTL3 in HSCs. We show that METTL3 negatively regulates the stability of GPR161 transcripts, which leads to decreased GPR161 expression through an m^6^A-dependent mechanism. Several studies have illustrated an important role for GPR161 during normal development and pathological process. An 8 bp mutation in GPR161 exhibits neural tube defects and congenital cataracts, due to a premature stop codon and truncated C terminus (Matteson, Desai et al., 2008). GPR161 knockdown in in the zebrafish embryo resulting in aberrant cardiac morphogenesis (Leung, Humbert et al., 2008). GPR161 localizes in primary cilia inhibits Shh signaling through Gαs-induced cyclic adenosine monophosphate (cAMP) accumulation, resulting in PKA-mediated processing of Gli3 (Shimada, Hwang et al., 2018). Moreover, GPR161 participates in a signaling network, characterized by GPCR activation and induction of adenyl cyclase–cAMP–PKA, that confers resistance to MAPK pathway inhibition in melanoma (Johannessen, Johnson et al., 2013). Overexpression of GPR161 in human triple-negative breast cancer correlated with poor prognosis (Feigin, Xue et al., 2014). Our further functional studies illustrated that GPR161 is an essential target of METTL3 in HF. As critical downstream targets of GPR161, Gli3 levels can also be indirectly regulated by METTL3 in HF. Through this axis, increased expression of METTL3 and decreased GPR161 expression in HF promotes HSC proliferation and inhibits its apoptosis. Of course, besides GPR161, other potential targets of METTL3 identified herein might also be important downstream targets of METTL3 and may partially mediate the overall function/effects of METTL3 in HF (and other pathological process), which warrants further systematic investigation.

Collectively, our results provide mechanistic insights into how expression of m^6^A modulators is regulated in HF and suggest a potential therapeutic strategy for selectively treating HF by targeting METTL3 GPR161/Gli3 axis. This work will inspire future explorations of the effects of post-transcriptional modifications on other RNA molecules.

## Materials and Methods

### Human liver tissues

The study protocols concerning human subjects were approved by the First Affiliated Hospital of Anhui Medical University (Permit No.20200734), and are consistent with the principles of the Declaration of Helsinki. The six normal liver tissues (without fibrosis) were obtained from transplant donors or patients with cholecystolithiasis who underwent cholecystectomy in the First Affiliated Hospital of Anhui Medical University. The 18 fibrotic and cirrhosis liver tissues were obtained from patients with HBV or HCV infection who underwent liver biopsy, or histologically diagnosed liver cirrhosis who underwent liver transplantation.

#### Fibrosis mouse model establishment

All experiments using mice were approved by the Institutional Animal Experimental Ethics Committee of Anhui Medical University (Permit No.LLSC20200730). Six-week-old male C57BL/6J mice (weight: 18-22 g) were purchased from the Laboratory Animal Center of Anhui Medical University. Care and maintenance of the animals was approved by the Institutional Animal Experimental Ethics Committee. All mice were housed in a temperature-controlled room (21℃ ± 1℃) under condition of 55% ± 10% humidity and a 12 hours light/dark cycle. Mice were placed in a pathogen-free animal facility with access to water and food, *ad libitum*.

To induce chronic liver fibrosis, six-week-old male mice were injected with CCl4 (Sigma-Aldrich, 2 mL/kg, in corn oil at a ratio of 1:4, i.p., twice a week for 4 weeks).

Animals were sacrificed 18 hours after the last injection of CCl4. Serum samples, liver tissues, and primary HSCs and primary HCs were collected for subsequent analyses.

For BDL model, the bile duct was doubly ligated with 6–0 silk. Sham-operated mouse were operated on similarly as control of C57BL/6 mice, except the bile duct was not ligated. Serum samples, liver tissues and primary HSCs were collected 21 days after the operation.

#### Recombinant adeno-associated virus-mediated HSC-specific METTL3 silencing and GPR161 overexpression in mice

The promoter of mouse *Lrat* gene (-1071∼ 0 relative to translation initiation site) was insert into the plasmid to mediated HSC-target silencing of METTL3. HSC-specific METTL3 silencing *in vivo* was established via liver in situ injection of recombinant adeno-associated-virus (Ad)-packed small hairpin RNA against METTL3 labeled with the luciferase protein (Ad-Lrat-shM3-Luc). The overexpression of GPR161 in HSCs was mediated by Ad-packed GPR161 overexpression plasmid labeled with enhanced Green Fluorescent Protein (Ad-Lrat-GPR161-eGFP). Mouse was slowly injected with 50 μL Ad-Lrat-shM3-Luc or Ad-Lrat-GPR161-eGFP at a concentration of 1 × 10^11^ v.g/mL/mouse using a 0.5 mL insulin syringe. One week later, mouse HF model was induced by injecting of CCl4 (2 ml/kg, CCl4 dissolved in corn oil at a ratio of 1: 4, i.p., biweekly, 4 weeks). The location of the virus accumulates in the mouse was indicated via an IVIS Lumina III Imaging System (Caliper Life Sciences, USA). Animals were sacrificed 18 hours after the last injection of CCl4. Serum samples, liver tissues, and primary liver cells were harvested for subsequent analyses.

#### Isolation of primary hepatic stellate cells and hepatocytes

A two-step collagenase perfusion procedure was used to isolate mouse hepatocytes as previously reported with slight modifications. Briefly, mice were perfused through the superior van cava by 1 x Hank’s balanced salt solution (HBSS) (GIBCO, 14175500BT) containing 15 mM HEPES (GIBCO, 15630080) and 1% penicillin-streptomycin (GIBCO, 15140163) (Step 1), followed by the collagenase digested buffer 1x HBSS containing 15 mM HEPES, 1% penicillin-streptomycin, pronase E (0.4 mg/mL) and collagenase IV (0.8 mg/mL, Thermo Fisher Scientific, 17104019) (Step 2). Liver was removed and digested *in vitro* with collagenase IV (0.5 mg/mL), pronase E (0.5 mg/mL), and DNAse I (0.02 mg/mL). The cell suspensions were filtered through a 70 μm nylon mesh, centrifuged at 50 x g for 5 minutes at 4 ℃, and the cell pellets (hepatocytes) were collected. The HSCs were separated using a Nycodenz (CAS no., 66108-95-0, I134719-25 g, Iohexol) gradient centrifugation. The HSC layer was treated with clodronate liposomes (5 mg/mL) for 4 hours at 37 °C to remove the contamination of macrophages as described (Inzaugarat, Johnson et al., 2019).

#### Histopathology, Sirius red and Immunostaining

Liver tissues were fixed, embedded in paraffin, and processed on slides for hematoxylin-eosin (H&E) or Sirius red staining. Histopathological staging was performed by an experienced pathologist. Primary antibodies used to perform immunostaining were: anti-α-SMA (bs-0189R, Bioss, 1:50 dilution), anti-METTL3 (ab195352, Abcam, 1:200 dilution), anti-GPR161 (ab58679, Abcam, 1:200 dilution). Following overnight incubation with primary antibodies, horseradish peroxidase (HRP) or Alexa-Fluor 488- or Alexa-Fluor 546-conjugated second antibody (Thermo Fisher Scientific) was applied for 1 hour. Mounting solution containing 4, 6-diamidino-2-phenylindole (DAPI) was used to counterstain the nuclei. The stained sections were scanned using Panoramic MIDI (3D HISTECH, Hungary) and viewed using CaseViewer slice software.

#### Serum analysis

Blood samples collected from mice were used to measure serum alanine aminotransferase (ALT) and aspartate aminotransferase (AST) levels. Serum ALT and AST levels were measured using alanine aminotransferase/aminotransferase assay Kits (C009-2-1/C010-2-1, Nanjing Jiancheng Bioengineering Institute, China) according to the manufacturer’s instructions. Absorbance at 510 nm was measured using the SpectraMax iD3 Multi-Mode Microplate Reader (Molecular Devices, USA).

#### Cell culture, treatment, and transfection

LX2 cells were cultured in the Dulbecco’s modified Eagle medium (DMEM, 11965092, Gibco, USA) containing 10% fetal bovine serum (FBS, 10099141, Gibco, USA) and cultured in a complete medium (CM-M041, Procell, China) at 37 °C in a humidified atmosphere containing 5% CO_2_. The LX2 cells and primary HSCs were activated with TGF-β1 (40 ng/mL, 100-21, Peprotech, USA) for 24 hours. Target gene knockdown or overexpression in primary HSCs was performed using the Lipofectamine™ 3000 Transfection Reagent (L3000015, ThermoFisher, USA). Information on the siRNAs used in this study has been listed and provided (**Table EV3**).

#### Establishment of stable knockdown and overexpression cells

Stable knockdown or overexpression of target genes in LX2 cells was achieved via lentiviral-packed short hairpin RNA (shRNA) or overexpression plasmid delivery. The transfected cells were selected with puromycin (1 μg/mL) before use. Information of the shRNAs sequences used in this study has been listed and provided (**Table EV3**).

#### Protein extraction and western blotting

Cells were subjected to washing steps twice with chilled phosphate-buffered saline (PBS) and were ruptured using the radioimmunoprecipitation (RIPA) buffer (P0013B, Beyotime, China) containing 50 mM Tris (pH 7.4), 150 mM NaCl, 1% Triton X-100, 1% sodium deoxycholate, 0.1% sodium dodecyl sulfate (SDS), 5 mM EDTA, PMSF, and phosphatase inhibitor cocktail. Cell extracts were centrifuged for 30 minutes at 12,000 x *g* and supernatants were collected. Nucleic and cytoplasmic proteins from LX2 cells were separated using the CelLytic™ NuCLEAR™ Extraction Kit (BB-3115, BestBio, China) following the manufacturer’s instructions. Protein concentrations were quantified using Omni-Easy™Instant BCA Protein Assay Kit (ZJ102, epizyme, China), and equal amounts (30-50 μg) of protein were separated by performing SDS-PAGE (10% or 12%) and then transferred onto polyvinylidene difluoride (PVDF) membranes (IPVH00005, Millipore, USA). Membranes were blocked for 1 hours with 5% skim milk at room temperature and then incubated overnight at 4 °C with the primary antibody. Subsequently, the membranes were washed and incubated for 1 hours with the appropriate secondary antibodies conjugated to horseradish peroxidase. Densitometry of the blots was detected using the Enhanced Chemiluminescence Kit HRP (SQ101, EpiZyme, China) in the ChemiDoc^TM^ MP Imaging System (Bio-Rad, USA). Information on the primary antibodies used has been listed and provided (**Table EV4**).

#### RNA extraction and quantitative real-time PCR

Total RNA from human patient samples, mouse liver and cell lines were exptracted using the TRIzol^TM^ reagent (15596018, Invitrogen, USA), and reverse transcribed into complementary DNA using Evo M-MLV RT Premix for qPCR (AG11706, Accurate Biology, China). The resulting cDNAs were used for PCR using SYBR® Green Premix Pro Taq HS qPCR Kit (AG11701, Accurate Biology, China). All PCR reactions were conducted using the Bio-Rad CFX-96 (Bio-Rad, USA). All data were normalized against endogenous GAPDH controls of each sample. Relative quantitation analysis of gene expression was performed using the 2^−ΔΔCt^ method. Information on the primer sequences has been listed and provided (**Table EV5**).

#### Cell cycle and apoptosis assays

For cell cycle analysis, transfected cells were collected and measured using the Cell Cycle and Apoptosis Analysis Kit (C1052, Beyotime, China) following the manufacturer’s instructions. Collected cells were fixed in 70% ethanol at 4 ℃ overnight, and then incubated with 25 μL PI and 10 μL ribonuclease A, incubated at 37 ℃ for 30 minutes and then applied to flow cytometer directly. Cell apoptosis assay was validated with FITC Annexin V Apoptosis Detection Kit I (556547, BD Biosciences) according to the manufacturers’ protocol. 1 x 10^6^ cells were collected and washed twice with cold PBS, and then resuspended in 100 μL 1 x binding buffer and adjust concentration to 1 x 10^6^ cells/mL. Add 5 μL of FITC Annexin V and 5 μL PI to the suspension, gently vortexed the cells, and incubated for 15 minutes at room temperature in the dark. Subsequently, add 400 μL of 1 x binding buffer to each sample and analyzed by flow cytometry within 1 hour. The samples were transferred onto ice before subjected to flow cytometry. Flow cytometry was performed with CytoFLEX flow cytometer (Beckman Coulter, USA), and the data were analyzed by CytExpert software (Beckman Coulter, USA).

#### EdU staining

EdU staining was conducted using the Cell-Light EdU Apollo567 In Vitro Kit (C10310-1, Ribio, China) according to the manufacturer’s introductions. Transfected LX2 cells were seeded on cover glasses and incubated for 24 hours, subjected to treatment with 50 μM EdU, and incubated for another 2 hours. Subsequently, the cells on cover slips were fixed with 4% paraformaldehyde at room temperature for 30 minutes, and then 0.5% Triton X-100 was added to permeabilize the cells. Subsequently, Hoechst 33342 was added to stain nuclear acids. Signals were detected using an inverted fluorescence microscope (Olympus IX83, Tokyo, Japan).

#### Migration assays

For conducting the scratch wound healing assay, parallel scratches were inflicted behind a six-well plate at a distance interval of 0.5-1 cm. Stable knockdown or overexpression cells were seeded into six-well plates and activated using TGF-β1 (40 ng/mL). When the cells reached approximately 90% confluency, scratches perpendicular to the parallel were inflicted using a sterile spearhead. After 12 hours of culture, images of cell motility occurring close to the scratches were captured using an inverted fluorescence microscope (Olympus IX83, Tokyo, Japan). The cell migration ability was reflected by the width of the scratch. Each experiment was repeated at least thrice.

For the transwell assay, 1 x 10^4^ METTL3-overexpressing or METTL3-silencing LX2 cells and the corresponding control cell in serum-free DMEM were added to each upper inset pre-coated with an 8.0-µm Pore Polycarbonate membrane. DMEM containing 10% FBS was added to the lower chambers. After 24 hours, the cells attached to the upper surface of the filter membranes were removed, and the migrated cells on the lower surface were stained with 0.5% crystal violet. The extent of migration was observed using an inverted fluorescence microscope (Olympus IX83, Japan).

#### MeRIP-Seq, RNA-Seq, and data analyses

m^6^A-specific methylated RNA immunoprecipitation using next-generation sequencing (MeRIP-Seq) was performed by LC-Bio-Technology Co., Ltd. (Hangzhou, China). Total RNA was isolated and purified using the TRIzol reagent following the manufacturer’s instructions. Approximately 25 μg of total RNA was used to deplete ribosomal RNA according to the Epicentre Ribo-Zero Gold Kit (MRZG126, Illumina, USA) manual. Following purification, the ribosomal-depleted RNA was fragmented into small pieces, and approximately 5 μg of the cleaved RNA fragments were incubated with anti-m^6^A-specific antibody (202003, Synaptic Systems, Germany) in IP buffer (50 mM Tris-HCl, 750 mM NaCl, and 0.5% Igepal CA-630) for 2 hours at 4 ℃. Then, the IP RNA was reverse-transcribed to generate cDNA ussing SuperScript™ II Reverse Transcriptase (1896649, Invitrogen, USA), which was then used to synthesize U-labeled second-stranded DNAs. Subsequently, the mixture was immunoprecipitated using Dynabeads® Protein A (10002D, ThermoFisher, USA) and rotated for an additional 2 hour at 4 ℃. After performing washing steps four times using IP buffer, the m^6^A IP portion was eluted from the beads using m^6^A-elution buffer and then extracted with the TRIzol reagent according to the manufacturer’s instructions. Purified RNAs were used to generate the RNA-Seq library. Both the input sample and the m^6^A IP portion were subjected to 150-bp paired-end sequencing using the Illumina Novaseq^TM^ 6000 following the manufacturer’s instructions.

Paired-end reads were harvested using the Illumina Novaseq 64000 sequencer, and subjected to quality control using Q30. After 3′ adaptor-trimming and low-quality reads were removed using the fastp software (https://github.com/OpenGene/fastp), the reads were aligned to the reference genome (mm10) using HISAT2 (http://daehwankimlab.github.io/hisat2). Mapped reads of IP and input libraries were provided for R package exomePeak (https://bioconductor.org/packages/exomePeak), which identifies m^6^A peaks in bed or bigwig format that can be adapted for visualization using the IGV software (http://www.igv.org). HOMER (http://homer.ucsd.edu/homer/motif) was used for exploring de novo motif, followed by localization of the motif with respect to peak summit. Differentially expressed RNAs were selected with |log2(fc)| ≥ 1 and *p* value < 0.05, using the R package edgeR (https://bioconductor.org/packages/edgeR).

Total RNA isolated from METTL3 silencing group in primary HSCs, and GPR161 overexpressed LX2 cells were used for RNA-Seq as described above.

#### Quantification of global m^6^A

Total RNA was isolated from different cells using the miRNase Mini Kit (217004, QIAGEN, Germany). Changes in the global m^6^A modification levels in mRNAs were determined using EpiQuik m^6^A RNA Methylation Quantification Kit (Colorimetric) (P-9005, Epigenetic, USA) following the manufacturer’s instructions. Poly-A-purified RNA (200 ng) was used for each sample analysis.

#### m^6^A dot blot assay

Total RNA was isolated from different cells using the miRNeasy Mini Kit (217004, QIAGEN, Germany) according to the manufacturer’s instructions and quantified using the DS-11 Sepectrophotometer (DeNovix, USA). For conducting the m^6^A dot blot assay, RNA samples were denatured at 70 ℃ for 5 minutes, and subsequently, equal amounts of serial-diluted mRNA were added to a nitrocellulose filter membrane. After UV crosslinking, the membranes were blocked with 5% skim milk and incubated with anti-m^6^A antibody (ab195352, Abcam, 1:1000 dilution) overnight at 4 ℃. Subsequently, the secondary antibody was incubated for 1 hour at room temperature. The membrane was exposed using the Enhanced Chemiluminescence Kit HRP (SQ101, EpiZyme, China) in the ChemiDoc^TM^ MP Imaging System (Bio-Rad, USA).

#### Chromatin immunoprecipitation (ChIP)-qPCR

Following transfection, LX2 cells were stimulated with TGF-β1 (40 ng/mL), and the corresponding control cells were cultured to ∼80%-90% confluency. Subsequently, ChIP assay was performed using the SimpleChIP^®^ Enzymatic Chromatin IP Kit (Magnetic Beads) (9003, Cell Signaling Technology, USA) according to the manufacturer’s instructions. The following antibodies were used to perform immunoprecipitation of the crosslinked protein-DNA complexes: rabbit anti-H3K27ac and normal rabbit IgG. The immunoprecipitated DNA was purified for performing RT-qPCR analysis.

#### RNA immunoprecipitation (RIP) assay

RNA immunoprecipitation (RIP) experiments were performed using the Magna RIP^TM^ RNA-Binding Protein Immunoprecipitation Kit (17-700, Millipore, USA) according to the manufacturer’s instructions. Briefly, LX2 cells were lysed using the RIP lysis buffer. Cell lysates were immunoprecipitated using the anti-METTL3 antibody (ab195352, Abcam, USA), anti-YTHDF2 antibody (ab220163, Abcam, USA), or control IgG antibody at 4℃ overnight, followed by RNA purification. Finally, immunoprecipitated RNA was analyzed using RT-qPCR.

#### RNA stability

To measure RNA stability in LX2 stable METTL3-silenced cells (shM3) and the corresponding control cells (shNC), 5 μg/mL of actinomycin D (Act-D, 15021, Cell Signaling Technology, USA) was added to the cells. After incubation at the indicated times, total RNAs was harvested, and used for RT-qPCR. The relative half-life (t_1/2_) of *GPR161* mRNA was estimated according to the methods described by previously published studies (Chen, Ezzeddine et al., 2008). When cultured with Act-D, mRNA transcription is turned off. The degradation rate of RNA (*K_decay_*) was calculated using the following equation:

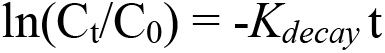

where t is represents the transcription inhibition time, and C_t_ and C_0_ represent the mRNA concentration at time t and time 0. The *K_decay_* can be derived by applying the exponential decay fitting of C_t_ /C_0_ versus time t. Therefore, the half-life (t_1/2_) (C_t_ /C_0_ = 1/2) can be calculated from the degradation rate as follows: t_1/2_ = ln2/*K_decay_*

#### Gene-specific MeRIP-qPCR

First, m^6^A immunoprecipitation (MeRIP) was performed using the EpiQuik™ CUT&RUN m^6^A RNA Enrichment Kit (P-9018, EPIGENTEK, USA) following the manufacturer’s instructions. Briefly, total RNA was extracted using the SteadyPure Universal RNA Extraction Kit (AG21017, Accurate Biology, China) with additional performance of DNase I on-column digestion. Then, 200-400 ng mRNA was saved as input sample, and approximately 10 μg total RNA was incubated with an immunocapture solution (immune capture buffer 174-189 μL, m^6^A antibody or non-immune IgG 2 μL, RNA sample 5-20 μL, Affinity beads 4 μL). The mixture subjected to rotation at room temperature for 90 minutes, and 10 μL of Nucleic digestion enhancer and, 2 μL of the cleavage enzyme mix were added to each tube and incubated at room temperature for 4 minutes. After performing washing steps three times using the wash buffer, the m^6^A IP portion was resuspended with 20 μL protein digestion solution (mixing proteinase K with protein digestion buffer at 1:10 dilution), followed by incubation at 55 °C for 15 minutes in a thermocycler (without a heated lid). The solution was then transferred to an unused PCR tube, followed by the addition off 20 μL RNA purification solution, 160 μL 100% ethanol, and 2 μL RNA-binding beads, and incubated for an additional 5 minutes at room temperature to facilitate the binding of RNA to the beads. Subsequently, the m^6^A-immunoprecipitated RNAs were released from the beads using 13 μL of elution buffer. The primer sequences were shown in **Table EV6**.

#### m^6^A methylation site prediction, plasmid construction, and luciferase reporter assay

Potential m^6^A sites were predicted using the online tool SRAMP (http://www.cuilab.cn/sramp/). The DNA fragments of GPR161-CDS and GPR161-3′ UTR regions containing the wild-type m^6^A motifs and mutant motifs (“A” in m^6^A motif was replaced by “G”) were directly synthesized by Hanbio Biotechnology Co., Ltd. (Shanghai, China). Wild-type and mutant GPR161-CDS and GPR161-3′ UTR were inserted downstream of the firefly luciferase of the pcDNA3.1 vector. METTL3 wild-type and mutant (C294A and C326A) plasmids and the corresponding control plasmid were constructed by Hanbio Biotechnology Co., Ltd. (Shanghai, China). For conducting dual-luciferase reporter assays, 200 ng wild-type or mutant GPR161-CDS (or GPR161-3′ UTR), 200 ng pcDNA3.1-METTL3 (or pcDNA3.1-METTL3-Mut or pcDNA3.1), and 20 ng pRL-TK (Renilla luciferase control reporter vector) were co-transfected into pri-HSCs in 24-well plates. The relative luciferase activities were assessed 48 hours post-transfection using the Dual-Luciferase Reporter Assay System (Promega, USA). Each experiment group was repeated in triplicate. Information on the primer sequences used for plasmid construction has been listed and provided (**Table EV7**).

##### (1) GPR161-CDS with wild-type m^6^A sites

CAAGGCCCGAAAGGTGCACTGTGGCACGGTGGTCACTGTGGAGGA**GG<u>A</u>CT**CTCAGAGGAGCGGGAG GAAGAATTCTAGTACCTCCACTTCCTCCTCCGGCAGTAGGAGGAATGCCCTTCAAGGAGTGGTCTATTCAGCTAACCAGTGCAAAGCCCTCATCACCATCCTGGTGGTCATTGGCGCCTTCATGGTCACCTGGGGCC CCTACATGGTTGTCATTACCTCAGAGGCACTCTGGGGGAA**GA<u>A</u>CT**GTGTCTCCCCAACCCTGGAGACT TGGGCCACATGGCTGTCCTTTACCAGTGCCATCTGCCACCCTCTGATCTAC**GG<u>A</u>CT**CTGGAACAAGAC TGTGCGCAAGGAGCTCCTGGGCATGTGCTTTGGGGACCGTTATTACCGGGAATCCTTTGTGCAGCGACAGAGGACCTCCAGGCTCTTCAGCATTTCCAACAGG

##### (2) GPR161-CDS with mutant m^6^A sites

CAAGGCCCGAAAGGTGCACTGTGGCACGGTGGTCACTGTGGAGGA**GG<u>G</u>CT**CTCAGAGGAGCGGGA GGAAGAATTCTAGTACCTCCACTTCCTCCTCCGGCAGTAGGAGGAATGCCCTTCAAGGAGTGGTCTAT TCAGCTAACCAGTGCAAAGCCCTCATCACCATCCTGGTGGTCATTGGCGCCTTCATGGTCACCTGGGG CCCCTACATGGTTGTCATTACCTCAGAGGCACTCTGGGGGAA**GA<u>G</u>CT**GTGTCTCCCCAACCCTGGAGA CTTGGGCCACATGGCTGTCCTTTACCAGTGCCATCTGCCACCCTCTGATCTAC**GG<u>G</u>CT**CTGGAACAAG ACTGTGCGCAAGGAGCTCCTGGGCATGTGCTTTGGGGACCGTTATTACCGGGAATCCTTTGTGCAGCGACAGAGGACCTCCAGGCTCTTCAGCATTTCCAACAGG

##### (3) GPR161-3′ UTR with wild-type m^6^A sites

AAGCTTCAGAGGGGAGCGGGGCCGCAGCGGAGCTACCG**GG<u>A</u>CT**GGTGCCTGTGCCAAGACCCTGGA ACATGTCACCACTCAGAGAGGGCCAGACGCCGGGAAGGAAACATCTGGTATGCAGCCAGGAGGATTGAAAATACTGTCTCCAGTCTCTGACTTCAGTTGTCTTCTGTAAAGCACATGTGGGCTGCTTCAGGGCTCCTGGGATGGAACGAGGCTTCGAAAGCACACGTTAGAAGCAGATGGAGATGTGGCTGGGGCATCAGGAGGACTAGAGATGGTGGATGTGAAAGGGAAGCCACCATGTGGCTCTTGTCTTCAATGGTATCCAGCTGCTTTAGCTGTCCCTGGTCTCAGTGAGGCCTCTAGAGGCAATCCCTACTCCGTGCCCCTCATGGGGACAGAACGGCTCACCCTAGCACTGTCAGACCATCTGGGGGCTCTGTGTAGTTGATCCAGGAAGCGTGGGACCACAAGTACTAGACTAAATGCAAAAACAGTCCAGAGAGCATCCGAGTAGAGCACCCCACATAGACACCC CATATCTATTGCCCTAGTTA**GG<u>A</u>CT**TGCTCTTGGGTAGCACCTGCTGCTGGCCAATAGAAACAGCCCAC AGGCCACCCAAGATGTGGCCACCCCACCCACTTCCTTGTTGACAGATTTTTATTATTACTATTATTATTTGAAAAGAAGGGAGAAACCACTATGAAAGACAAGGGCGAATACCAAGCACTTCATTTCCAGAATAATAAAACCAGGAGGTTAGGGCATATCACTAAGCCTGGGACTTGAGGTCATACGCAGATGAGGGGAACTTG GGGGGAAATCCCTCACACGGGAGATATAGGAGATGCTAGAG**TG<u>A</u>CT**GGCAAACATAGCTGGCCTCAC TTTCCAGAG

##### (4) GPR161-3′ UTR with mutant m^6^A sites

AAGCTTCAGAGGGGAGCGGGGCCGCAGCGGAGCTACCG**GG<u>G</u>CT**GGTGCCTGTGCCAAGACCCTGGA ACATGTCACCACTCAGAGAGGGCCAGACGCCGGGAAGGAAACATCTGGTATGCAGCCAGGAGGATTGAAAATACTGTCTCCAGTCTCTGACTTCAGTTGTCTTCTGTAAAGCACATGTGGGCTGCTTCAGGGCTCCTGGGATGGAACGAGGCTTCGAAAGCACACGTTAGAAGCAGATGGAGATGTGGCTGGGGCATCAGGAGGACTAGAGATGGTGGATGTGAAAGGGAAGCCACCATGTGGCTCTTGTCTTCAATGGTATCCAGCTGCTTTAGCTGTCCCTGGTCTCAGTGAGGCCTCTAGAGGCAATCCCTACTCCGTGCCCCTCATGGGGACAGAACGGCTCACCCTAGCACTGTCAGACCATCTGGGGGCTCTGTGTAGTTGATCCAGGAAGCGTGGGACCACAAGTACTAGACTAAATGCAAAAACAGTCCAGAGAGCATCCGAGTAGAGCACCCCACATAGACACCC CATATCTATTGCCCTAGTTA**GG<u>G</u>CT**TGCTCTTGGGTAGCACCTGCTGCTGGCCAATAGAAACAGCCCAC AGGCCACCCAAGATGTGGCCACCCCACCCACTTCCTTGTTGACAGATTTTTATTATTACTATTATTATTT GAAAAGAAGGGAGAAACCACTATGAAAGACAAGGGCGAATACCAAGCACTTCATTTCCAGAATAATAAAACCAGGAGGTTAGGGCATATCACTAAGCCTGGGACTTGAGGTCATACGCAGATGAGGGGAACTTG GGGGGAAATCCCTCACACGGGAGATATAGGAGATGCTAGAG**TG<u>G</u>CT**GGCAAACATAGCTGGCCTCAC TTTCCAGAG

### Statistical analysis

All experiments were replicated at least thrice and the results are presented as mean ± SEM. Results that were normally distributed (p > 0.05 from Kolmogorov-Smirnov test) were compared with parametric statistical procedures (Student t test and ANOVA followed by Bonferron’s test for multiple comparisons). Non-normally distributed results were compared with non-parametric tests (Kruskall-Wallis one-way ANOVA and Mann-Whitney-U test). Significance was accepted at p < 0.05.

## Data availability

No data was uploaded to the public database. All the data were available upon rational request.

## Acknowledgements

This project was supported by the National Natural Science Foundation of China (82070628, U19A2001), the Universities Natural Science Foundation of Anhui Province (KJ2019A0233), the University Synergy Innovation Program of Anhui Province (GXXT-2019-045, GXXT-2020-063, GXXT-2020-025), and the Research Fund of Anhui Institute of Translational Medicine (2021 zhyx-B06). We apologize to colleagues whose work could not be cited owing to space constraints.

## Conflict of Interest

The authors declare that the research was conducted in the absence of any commercial or financial relationships that could be construed as a potential conflict of interest.

## Author Contributions

X.Y.P conceived and carried out experiments, analyzed the data, and wrote the manuscript. Y.H.B helped to analyze the data, and modified the manuscript. M.C, Z.Z.Q, L.W and H.M.Y carried out experiments. L.L and Z.H.Z provided human liver tissues. X.M.M and C.H participated in the design of the study. J.L. conceived the study and revised the manuscript. All authors read and approved the final manuscript.

